# Rescuing Troves of Data to Tackle Emerging Mosquito-Borne Diseases

**DOI:** 10.1101/096875

**Authors:** Samuel S.C. Rund, Micaela Elvira Martinez

## Abstract

According to the World Health Organization, every year more than a billion people are infected with vector-borne diseases worldwide. There are no vaccines for most vector-borne diseases. Vector control, therefore, is often the only way to prevent outbreaks. Despite the major impact of vectors on human health, knowledge gaps exist regarding their natural population dynamics. Even the most basic information—such as spatiotemporal abundance— is not available. Mosquitoes transmit malaria and the viruses causing Yellow Fever, West Nile, Dengue, Chikungunya, and Zika in the Americas. The Americas have a long history of mosquito control efforts, including the unsustained but successful *Aedes aegypti* eradication initiative. In the US, municipalities have independently created agencies for mosquito control and monitoring. We propose that the ensemble of US mosquito control agencies can, and should, be used to develop a national—and potentially international—system for Cross-Scale Vector Monitoring and Control (CSVMaC), in which local level monitoring and control efforts are cross-linked by unified real-time data streaming to build the data capital needed to gain a mechanistic understanding of vector population dynamics. Vectors, and the pathogens they transmit, know no jurisdictions. The vision of CSVMaC is, therefore, to provide data for (i) the general study of mosquito ecology and (ii) to inform vector control during epidemics/outbreaks that impact multiple jurisdictions (i.e., counties, states, etc.). We reveal >1000 mosquito control agencies in the US with enormous troves of data that are hidden among many data silos. For CSVMaC, we propose the creation of a nationally-coordinated open-access database to collate mosquito data. The database would provide scientific and public health communities with highly resolved spatiotemporal data on arboviral disease vectors, empowering new interventions and insights while leveraging pre-existing human efforts, operational infrastructure, and investments already funded by taxpayers.

## Mosquito Control & Surveillance

Mosquito transmitted infectious diseases pose a threat to public health worldwide. Over the past century, municipalities across the US have periodically launched mosquito control agencies to tackle existing or emerging disease threats, including the *Aedes aegypti* eradication initiative in the Americas (1947–1970), the malaria eradication program in the southern US (1947–1951), and the efforts to control WNV after its 1999 introduction into the US. The US CDC has guidelines for standardized and repeated collection of arboviral disease-vectors. Mosquito control agencies, thus, perform routine surveillance by trapping mosquitoes to measure their population size and inform their local efforts^*1, 2*^. Importantly, the primary role of mosquito control agencies is to locally mitigate vector populations; they carefully trap and taxonomically identify mosquitoes to monitor temporal changes in abundance. These agencies, therefore, generate long-term ecological time series of mosquito abundance, which collectively represent unprecedented ecological data that are spatially, temporally, and taxonomically rich. In addition, some agencies also perform in-depth phenotyping for insecticide resistance and arbovirus infection. We propose local level monitoring and control efforts should be cross-linked by unified real-time data streaming to create a national surveillance system for Cross-Scale Vector Monitoring and Control (CSVMaC). If the US were to create such a system, it could set the precedent for expanding such efforts to other disease vectors and being adopted by other countries that have vector control efforts operating within local jurisdictions.

In the US, clinical cases of mosquito-borne infectious diseases are nationally notifiable. There is state-level reporting to the CDC National Notifiable Disease Surveillance System (NNDSS) and summary tables are published in the Morbidity and Mortality Weekly Report (MMWR)^3^. CSVMaC in its essence is complementary to—and was inspired by—the NNDSS. A surveillance system for CSVMaC would need to be: standardized, centralized, and routine. This system would warehouse the vast amount of mosquito surveillance data that have been collected for decades and would provide the digital infrastructure for future weekly/monthly real-time data streaming. Here we (i) identify mosquito control agencies that could cross-link their data, revealing the potential breadth of CSVMaC, (ii) identify existing digital infrastructure that could be leveraged for national surveillance and (iii) demonstrate data can be cross-linked by collating those available from 14% of the agencies identified.

## The Tip of the Iceberg of Rich Ecological Data

We found 1054 mosquito control agencies (Table S1) scattered throughout the contiguous US (Fig 1), broadly defining a control agency as the local government authority responsible for mosquito control and surveillance. We attempted to locate an online presence for each agency. A total of 152 agencies had publically available data (i.e., live weekly/monthly updates and/or archived data). We collated all publicly available data going back to 2009, which included mine-able formats such as tables, graphs, spreadsheets, or GIS maps. Importantly, we only collated data collected using fixed mosquito traps. Fixed traps are in a fixed location and repeatedly sampled; their data, therefore, represent geo-tagged time-series.

**Fig 1.**
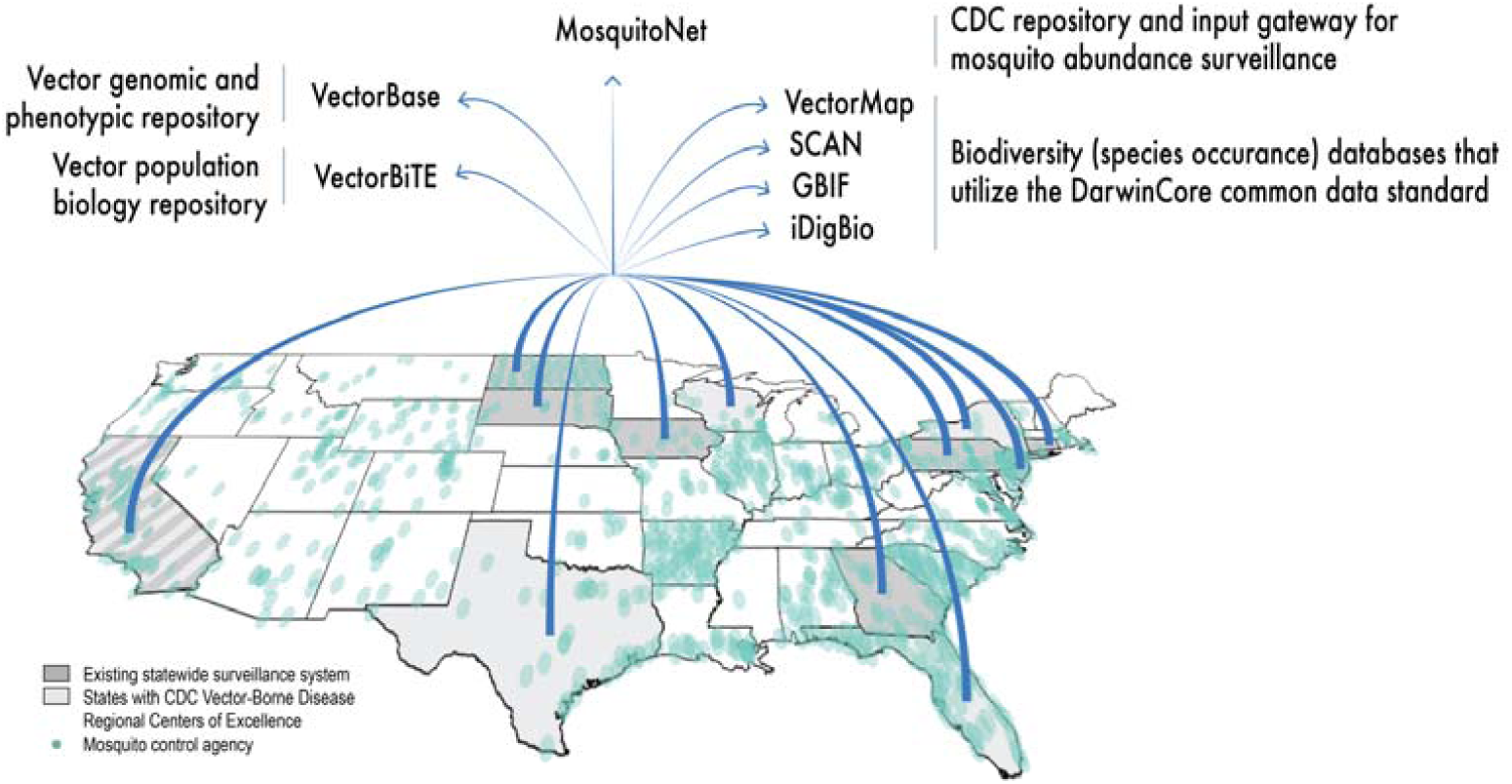
Existing infrastructure for nationwide mosquito surveillance. Points on map shows the location of the mosquito control agencies we identified. Dark grey states are those that have existing state-wide surveillance systems. These include California (CalSurv), North Dakota (North Dakota Department of Public Health), South Dakota (Mosquito Information Systems), Pennsylvania (Pennsylvania Dept. of Environmental Protection, Division of Vector Management), Iowa (Iowa State University, Department of Entomology), Connecticut (The Connecticut Agricultural Experiment Station), Georgia (Georgia Department of Public Health) and New Jersey (New Jersey Agricultural Experiment Station). Light grey states have newly funded CDC Vector-Borne Disease Regional Centers of Excellence. These are based at the University of Texas Medical Branch at Galveston, University of Florida, University of Wisconsin Madison, Cornell University, and University of California. Existing digital infrastructure that could be modified to serve as the National Vector Surveillance System (i.e., MosquitoNet) and/or be linked to the surveillance system (VectorBase, VectorBiTE, etc.) are shown.

A total of 148 agencies (14%) across 20 states had live—2016—or archived data. Iowa, Connecticut, New Jersey, and North Dakota have statewide open-access weekly surveillance dating back to 1969, 1998, 2003, and 2006, respectively; while, Pennsylvania, California, and South Dakota have state-wide systems that are not publically available. Some agencies provided trap-level data, while others reported mosquito counts aggregated from a geographic area; for example, a park, jurisdiction, or an ecological region.

In total, our dataset contained reports from > 600 unique agency-defined locations, > 39,000 instances when traps were checked, approx. 200,000 records, and documented > 15 million individual mosquitoes (Table S2). Records were each time-stamped with a collection date and geo-referenced. Taxonomic identification and resolution of reporting varied among districts. Often, mosquito reports were at the genus level; but 56 data sources reported species for at least a subset of mosquitoes, reflecting local knowledge of, and interest in, disease vector and nuisance species. With over 15 million mosquitoes trapped and identified by biologists on the ground, these data (Fig S1) represent a great deal of human effort. From these data we were able to reconstruct spatiotemporal abundance patterns across jurisdictions Fig S2.

Importantly, the data we collated are merely the tip of the iceberg. We focused on data from 2009 onward and only captured the small fraction of extant data that were available online in a mineable format. Large historical collections exist, with many agencies possessing finer-scale and longer-term data than publicly accessible. The majority of data remain hidden because there is no centralized repository and most agencies do not maintain webpages for reporting.

Fig 1 highlights the high resolution of mosquito monitoring. Monitoring, however, may be much more expansive than we observed. US military installations have conducted mosquito surveillance dating back to 1947^*4,5*^. In addition to mosquito abundance, which was the focus of this study, the proposed CSVMaC surveillance system could be made to include data on insecticide resistance and arbovirus-presence in mosquitoes. ArboNet, which is run by the US CDC and state health departments, is a national arboviral surveillance system for warehousing data complementary to ours^6^. It includes clinical cases of arbovirus infections in humans, mosquito testing for viruses, and sentinel animal reports. Unfortunately, ArboNet does not provide open-access live or high-spatiotemporal-resolution data.

## Surveillance Network Design

Our proposed national surveillance system would facilitate the flow of historical and ongoing mosquito abundance data into a centralized system at weekly or monthly intervals. CDC guidelines are already in place for standardized and repeated collection of vectors, to which data-reporting guidelines could be added. For national surveillance, it will be important to standardize what is reported and the units for reporting. In our collation efforts, we found that the format of existing data varied widely among agencies; however, agencies could standardize reporting without significant changes in their operational protocols. We propose agencies report the number of mosquitoes per trap—indicating if they counted females only, or both sexes—along with: trap type, location, attractants used, genus/species, date, the timeframe of collection, and any metadata necessary for data interpretation and integration.

Significant amounts of operational (i.e., on the ground) and digital infrastructure for CSVMaC already exist. There is, thus, no need to build a surveillance system from scratch. Existing digital infrastructure (Table S3) that could be modified for national surveillance includes that of VectorBiTE—a research coordination network funded by the US and UK, which integrates studies of vector behavior with transmission ecology—and VectorBase, the US National Institute of Allergy and Infectious Diseases Bioinformatics Resource Center, which warehouses genomic and phenotypic data from studies of vectors sampled worldwide. There are also biodiversity databases housing species occurrence records. In order to facilitate sharing, these databases utilize data standards known as Darwin Core^*7*^ and include generalist databases GBIF and iDigBio, SCAN (arthropods), and VectorMap (including US military mosquito records). We propose these, and other, existing digital systems should be actively threaded together. For national surveillance of the type we are proposing, a logical way forward would be to first have the multiple existing statewide mosquito surveillance systems come together; while the CDC and the five newly-funded CDC Centers for Excellence in Vector Borne Diseases, spearhead further data collation across the remaining US states. Once the streaming of abundance data is established, then the national surveillance system could be cross-linked with the data from other platforms, i.e.,VectorBiTE, VectorBase, etc.

As previously mentioned, many states report on mosquito viral presence/absence to ArboNet. This means local control agencies already have channels for streaming data to their state, which may be co-opted for reporting abundance data. Each state could subsequently stream records to the national system. In the future, the logistics of data streaming may be simplified with automated data collection. Next-generation traps—such as those from Microsofts’ Project Premonition and Biogents’ BG-Counter— connected to cellular networks, could automatically deliver data to the national surveillance system.

## Unrealized Potential

Mathematical models are relied upon to infer the risk of mosquito-transmitted infectious diseases. It is not possible to monitor individual mosquito-to-human or human-to-mosquito transmission events; therefore, transmission rates must be inferred by fitting transmission models to data. Fig 2 depicts the major transmission pathways in mosquito-borne disease systems. The notification of clinical cases of vector-borne diseases allows for the partial observation of transmission events, but without vector surveillance, we are missing the opportunity to observe a key component of the system: the mosquito population dynamics. Data, both historical and contemporary, from geographically independent regions are critical for model calibration and validation^*8–10*^; however, these data are lacking for arboviral disease vectors. Acheson and Kerr (2015) reviewed 29 vector-borne disease modeling publications, and revealed the lack of mosquito data. At least six of the publications they reviewed explicitly claimed insufficient data for model validation^*10*^. During the 2016 Zika outbreak, this data gap was particularly acute^*8, 11*^.

**Fig 2.**
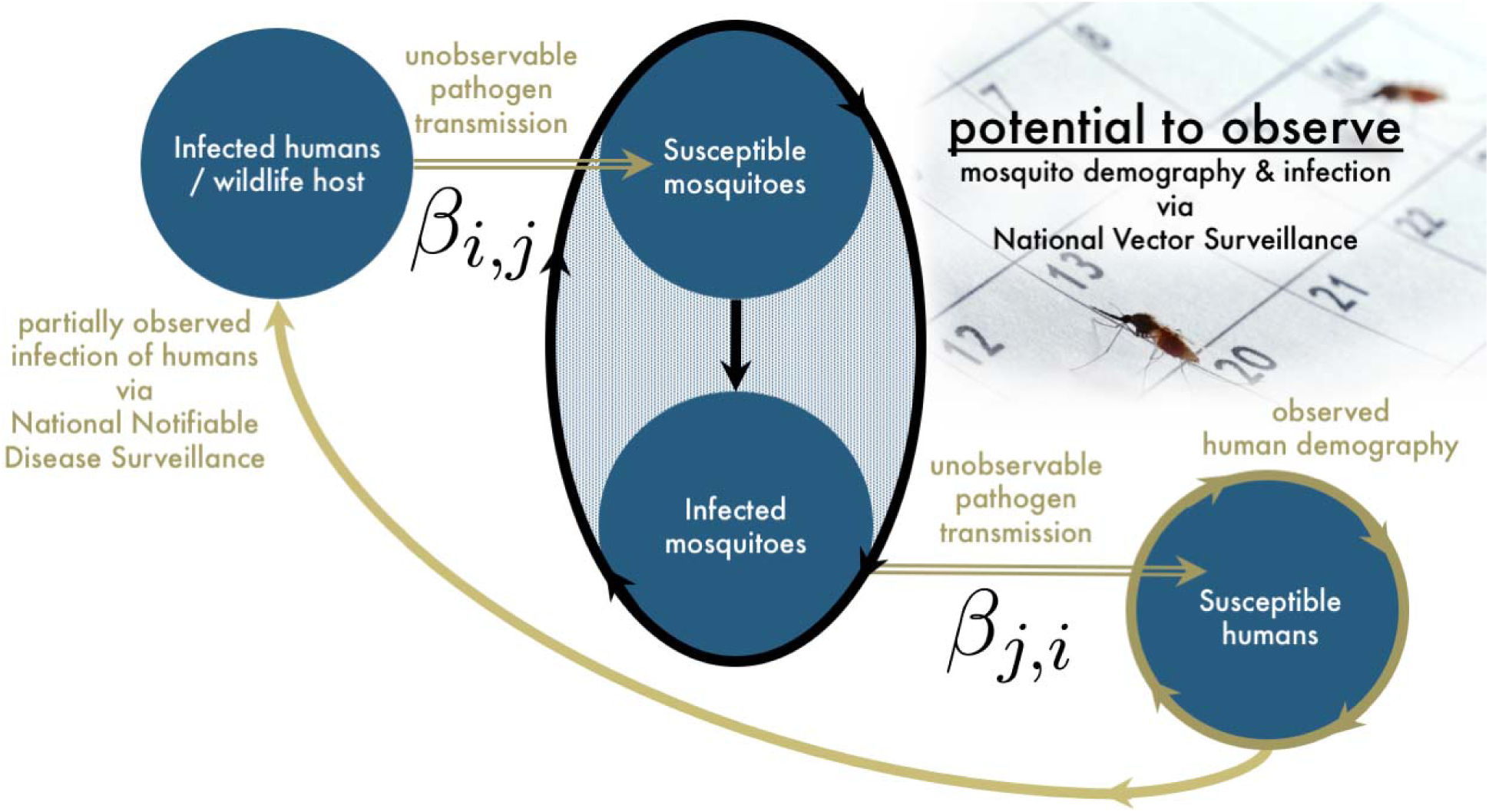
Generalized model schematic for mosquito-transmitted infections. Susceptible mosquitoes acquire infection by taking blood-meal from infected humans (e.g., for dengue or Zika virus) or wildlife (e.g., for West Nile virus). Infected mosquitoes may then transmit to susceptible humans. The two types of transmission events, human/wildlife-to-mosquito and mosquito-to-human, are not observable. However, these transmission rates, indicated by β_i,j_ and β_j,i_, respectively, can be inferred using mathematical transmission models coupled with data relating components of the system. Currently, the human piece of the data puzzle is in place, but mosquito data are lacking. Human demography data are readily available in the US, and infections in humans are partially observed via National Notifiable Disease Surveillance. Human infections are only partially observed because not all infections result in clinical diseases, and not all clinical cases are reported. The National Vector Surveillance System proposed here would fill in the mosquito data gap by providing mosquito demography (i.e., abundance) data that can be linked to arbovirus-presence in mosquitoes.

Rich longitudinal data of mosquito abundance would facilitate new analyses relating to mosquito biology and pathogen ecology that could inform the control of vectors during epidemics/outbreaks; especially those that impact multiple jurisdictions. It is important to note that mosquito trapping does not provide measures of absolute abundance of vectors. It can, however, be used to infer relative abundance. Data from existing mosquito control agencies represents the breadth of urban and ecological conditions in the US, and could be used to gain mechanistic insight^*12, 13*^ into the factors regulating vector populations and the pathogens they carry. Specifically, CSVMaC will allow for the study of:

1. variation in the relative abundance of vectors (i.e., seasonal variation within a geographic location and variation among geographic locations),
2. the mechanisms by which vector populations are regulated (e.g., how temperature or competition impacts vector abundance), and
3. local and metapopulation transmission dynamics of vector-borne pathogens.

The ability to quantify the relative abundance of vectors throughout the year and across the landscape would be powerful for understanding disease transmission. This is because spatially and temporally resolved relative abundance could be used to identify the window containing the high season for virus transmission within a geographic location, and how this window varies across the landscape. This information—when combined with (i) mechanistic transmission models, (ii) data on clinical disease (publicly available through the CDC), and (iii) human demography data—could be used to reveal pathogen metapopulation dynamics and inform disease control and eradication strategies. See Table S3 for clinical and demography data sources.

As for mosquito population biology, the observed spatiotemporal variation in vector abundance can be used to study the climatic and ecological factors regulating vector populations. For instance, vector abundance time series may be combined with other sources of big data, such as land use and meteorological data (Table S3), to determine how climate and vegetation work to shape vector abundance. A mechanistic understanding of how climate and ecological conditions impact vector populations is particularly important under climate change predictions and changes in urbanization^*14*^.

## The Way Forward

The increasing interconnectedness of human populations, global climate change, and the emergence of new vector-borne diseases necessitates diligent vector control that goes beyond jurisdictional boundaries. CSVMaC would be a simple low-cost solution for empowering foundational research on mosquito and infectious disease ecology. The CDC, being a federal organization charged with human health, and already housing the NNDSS and MMWR, is the obvious choice for facilitating CSVMaC. A CDC-based surveillance system would be highly visible internationally and therefore most likely to be used as a template for systems in other countries. The National Association of County and City Health Officials (NACCHO) and/or American Mosquito Control Association (AMCA), however, could also lead in coordinating this effort. Importantly, the CDC has created a repository for vector surveillance data called MosquitoNet.

MosquitoNet could serve as a gateway for mosquito agencies to enter data (particularly in areas not served by other state and regional infrastructure) as well as a final repository for the data. In other words, MosquitoNet could be the central digital infrastructure of the US National Vector Surveillance System. Disconcertingly, at present, MosquitoNet is not open access. We do not know why this is the case. This seems to go against the Health and Human Services principles on public access^*15*^. Mosquito surveillance is currently conducted with (i) tremendous tax-payer cost and (ii) large amounts human labor and expert knowledge, which has been ongoing for decades. Valid data standardization concerns exist, including the need to account for non-standardized collection protocols, but we call on the greater mosquito control and public health communities, as well as the US federal government, to maximize health and research benefits of mosquito control efforts by creating a US national vector surveillance system.

## Acknowledgments

This work was possible due to the efforts of the 148 mosquito control agencies who collected mosquito population abundance data and made their data available online. We thank research students Nathan Nixon, Ben Weise, Ben Chan, Jason Cleland, Ollie Lee, and Emma Stotter for their work in locating and digitizing data. Micaela Martinez was funded by a US National Science Foundation Postdoctoral Fellowship in Biology Award Number 1523757, and is currently funded by the NIH Directors Early Independence Award. Research reported in this publication was supported by the Office of the Director, National Institutes of Health, under Award Number DP5OD023l00. Samuel S.C. Rund was funded by the Royal Society (NF140517).

## Supplemental material

**Figure S1.**
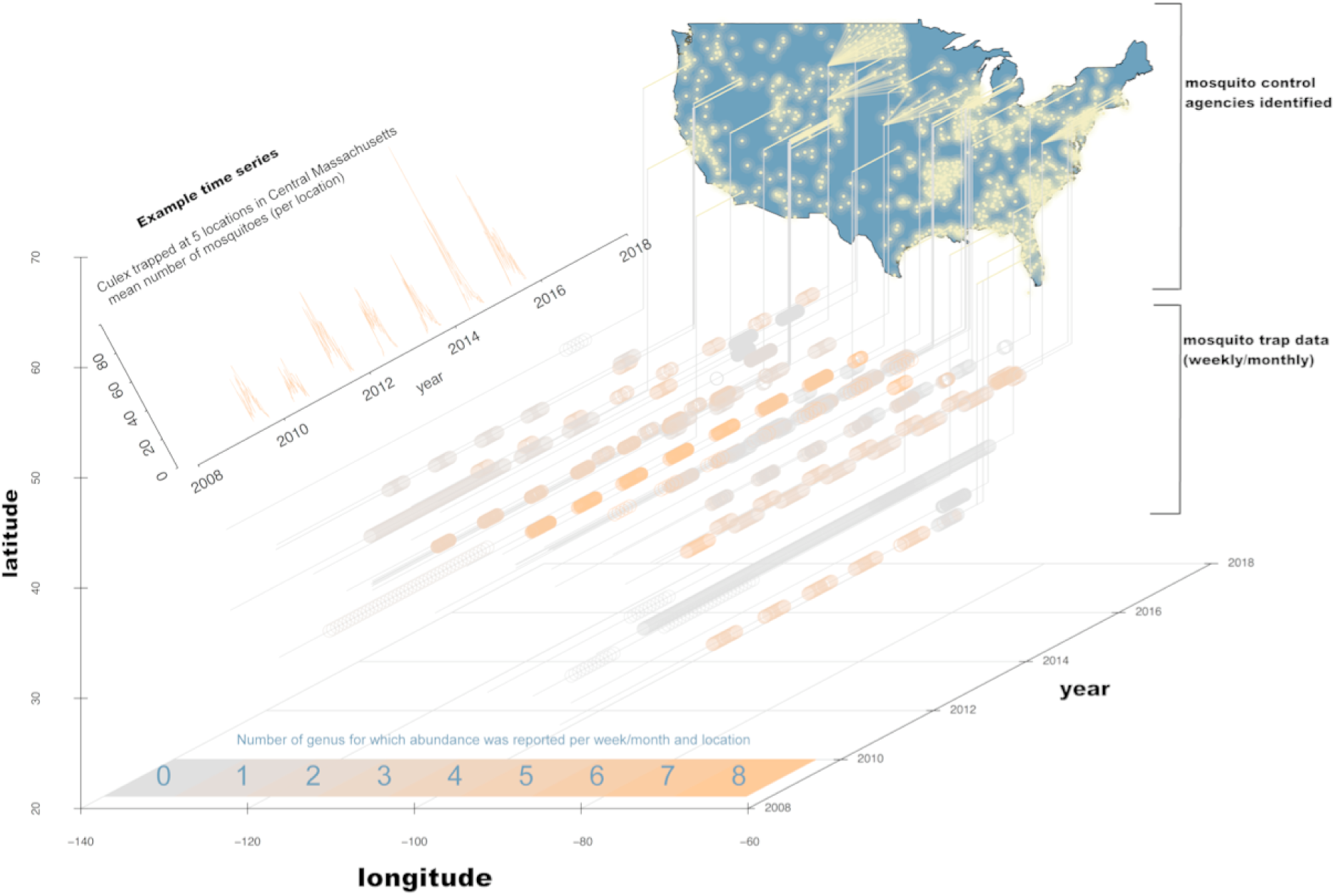
Map indicating the location of agencies we identified (yellow points). Some agencies made their data publicly available (those with a yellow line projection). Time series (grey and orange circles) show the collection dates and the number of genus whose abundance was reported in our data. Each time series represents one data silo. Despite the enormous amount of data (shown are > 39,000 trap collections), far more exist. Each agency is a potential data source. (inset) Example time series of Culex abundance from one agency, the Central Massachusetts Mosquito Control Project, which had five trap locations in their jurisdiction. Culex is of interest because it is the genus containing species that transmit WNV.

**Figure S2.**
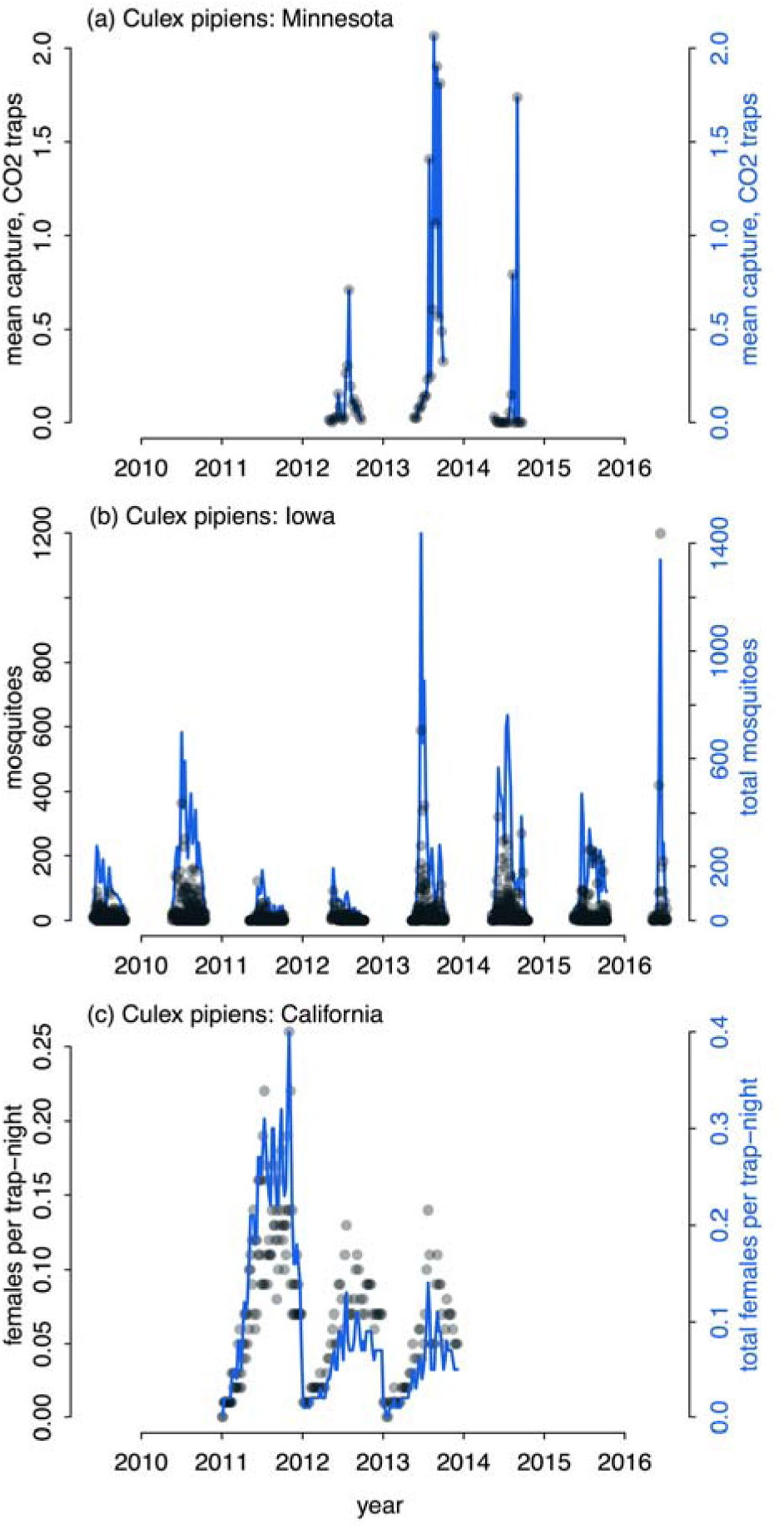
*Culex pipiens* seasonality from disparate regions of the country can be reconstructed from diverse data silos, despite differences in trapping and reporting protocols (e.g. ‘females per trap-night’, ‘mosquitoes,’ and ‘mean capture – CO_2_ traps’). Black points show the raw data from each trap within the state and correspond to the left y-axis. Blue time series are aggregated data across traps within the state, corresponding to the right y-axis. For any given taxonomic group, seasonal phenology may vary geographically due to variation in environmental conditions. We searched the compiled data for a taxonomic group for which we had 3+ years of data from multiple states. Data from *Culex pipiens* in Minnesota, Iowa, and California fit these criteria and were of particular interest because *C. pipiens* is the vector of West Nile Virus. We are able to reconstruct that the *C. pipiens* season was restricted to a narrow time frame (late summer) in Minnesota (a), the most northern of the three states. In Iowa and California, the *C. pipiens* season extended later into the fall (b-c). In California, the *C. pipiens* season began in the early spring. This demonstrates it is possible to unify mosquito abundance data collected by different people, using different collection protocols, and different reporting protocols.

